# A synthetic bacterium that degrades and assimilates poly(ethylene terephthalate)

**DOI:** 10.1101/2025.09.28.673679

**Authors:** Dekel Freund, Kesava Phaneendra Cherukuri, Raul Mireles, Joseph Kippen, Maya Shossel, Lianet Noda-García

## Abstract

Polyethylene terephthalate (PET) is the fourth most commonly used plastic worldwide. Like all plastics, post-consumer PET is poorly managed and accumulates in the environment, posing significant ecological threats. After 70 years of accumulation, microorganisms capable of degrading and assimilating PET have been isolated, demonstrating that PET can be broken down and converted into valuable cellular biomass or metabolic products. These natural isolates, however, are poorly characterized and challenging to genetically manipulate, which limits their further optimization and applicability. Here, we engineer a well-established synthetic biology chassis for the biodegradation and assimilation of PET. We modified the bacterium *Pseudomonas putida* KT2440 to heterologously express an active PET-hydrolytic enzyme extracellularly and to metabolize PET biodegradation products. The resulting strain, named PETBuster, was capable of growing on PET as the sole carbon source on solid and liquid media. We achieved 91% PET degradation after 21 days of culture, with a doubling time of 3.6 days, under mesophilic conditions. In this way, we demonstrate that PET fermentation is feasible, opening the door to the production of valuable chemicals from waste.

## Introduction

Plastics have become an integral part of modern life due to their durability, flexibility, and low production cost. Despite being introduced only about 70 years ago^1^, plastics are now found across a wide range of environments—including within living organisms^2^, in oceans^3^, landfills^4^, and even in some of the most remote and extreme places on Earth, such as the Mariana Trench^5^ and Mount Everest^6^. Global plastic production now exceeds 400 million metric tons annually and is projected to reach 700 million metric tons by 2030, equivalent to approximately 80 kilograms of plastic produced per person worldwide^7^. Without effective recycling solutions, the environmental impact is expected to intensify^8^. Polyethylene terephthalate (PET), the fourth most common plastic polymer^9,10^, consists of two fossil-derived monomers, ethylene glycol (EG) and terephthalic acid (TPA), linked by ester bonds^11^. PET is widely used in food packaging, beverage bottles, and synthetic fibers -in the U.S., PET bottled water accounts for 70.7% of all bottled water plastic volume^12^. In 2024, global PET production surpassed 82 million metric tons^13^.

PET is an ester, like other abundant biomolecules such as triglycerides and cutin^14^. Although esterases abound in biological systems^15^, only three natural bacterial isolates capable of degrading and assimilating PET have been reported (*Ideonella sakaiensis* 201-F6^16^, *Rhodococcus pyridinivorans* P23^17^, and *Pseudomonas chengduensis* BC1815^18^. These strains express one or more PET-hydrolyzing esterases (PETases) and can assimilate the monomers that result from PET biodegradation, TPA, EG, or both. However, they are not amenable to standard genetic engineering tools, which limits their optimization for different industrial settings and broader applications.

Motivated by these challenges, recent global efforts have focused on developing a synthetic, optimizable, and genetically tractable chassis for the degradation and assimilation of PET^19–21^. Such systems would serve as whole-cell biocatalysts for PET treatment, enabling processes in which PET is used as a feedstock for producing valuable biomass or compounds. Alternatively, they could be applied as high-throughput screening platforms to identify and resolve bottlenecks in PET fermentation, such as the development of faster PETases^22,23^.

Synthetic biology-friendly bacterial chassis, such as *Pseudomonas putida* and *Escherichia coli*, have been successfully engineered to have optimized TPA^24^ and/or EG metabolism^25–27^. It has been demostrated that valuable chemical compounds can be produced from them^20,28^. Moreover, in these and other engineerable microorganisms, it has been shown that PETases can be heterogeneously expressed extracellularly^19,20,29,30^. The combination of these systems—EG or TPA metabolism plus active PETases—has been attempted. *P. putida* KT2440, hereafter Pp_KT2440, was successfully engineered to assimilate Bis(2-Hydroxyethyl) terephthalate (BHET), a small PET oligomer with one TPA and two EG molecules containing two ester bonds^20^; and *Pseudomonas umsongensis* GO16 can degrade and assimilate PET polymers when other carbon sources are provided^21^. The use of the PET polymer as the sole carbon source for fermentation processes, however, has remained elusive.

Here, we report the successful engineering of Pp_KT2440 to grow on PET as the sole carbon source. By testing different system components, we identified a combination that integrates TPA metabolism with active extracellular PETase expression, establishing a fully synthetic bacterial platform capable of degrading and assimilating PET (Figure 1). Our strain, PETBuster, combines the native TPA metabolism of *P. umsongensis* GO16^31^ with an active EG pathway and the expression of the highly efficient engineered FAST_PETase^32^. As a result, PETBuster was able to degrade 91 ± 10% of PET within 21 days, with an average doubling time of 3.6 ± 0.8 days when using PET as the sole carbon source. Throughout the growth period, no accumulation of PET monomers was observed in the medium, indicating efficient uptake and metabolism of degradation products. This synthetic bacterium represents a significant step toward the practical application of a whole-cell biocatalyst for addressing one of the most common pollutants on Earth today.

**Figure 1.**
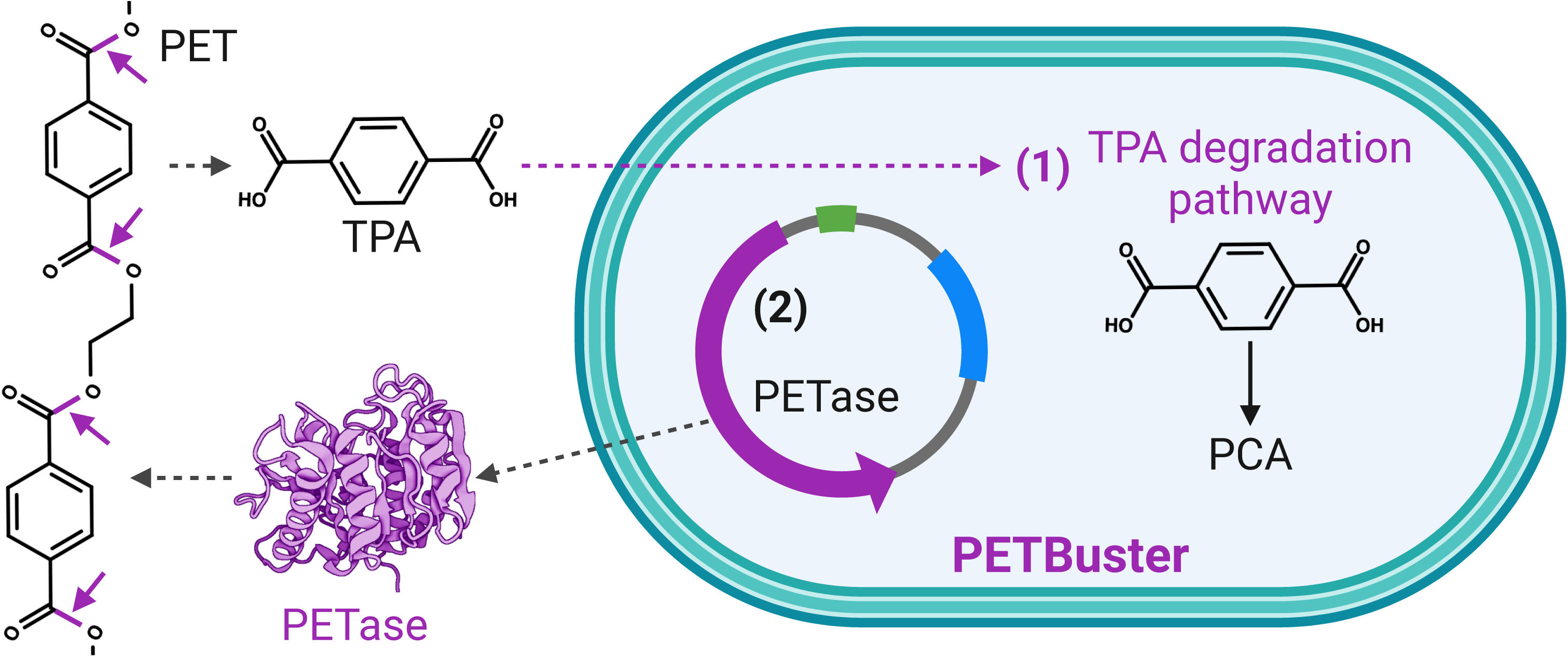
Schematic representation of the engineering strategy to create PETBuster. (1) The TPA catabolic pathway from *P. umsongensis* GO16, consisting of four enzymes, one transporter, and one transcriptional regulator (detailed in Figure 2), was integrated into the genome to enable TPA metabolism. (2) A plasmid was introduced to express and secrete an active extracellular PETase (detailed in Figure 3). The ester bonds hydrolyzed by PETase are indicated.

## Results

### Engineering the intracellular metabolism of PET degradation products

For Pp_KT2440 to degrade and use PET as its sole carbon source, the bacterium must be capable of metabolizing its monomers, EG or TPA. In Pp_KT2440, EG metabolism is available, although commonly silent ^24,25,27,30,33,34^. Moreover, studies have shown that in natural PET-eating bacteria, *I. sakaiensis* 201-F6^16^, *R. pyridinivorans* P23^17^, or engineered strains capable of surviving on BHET^20^, TPA is preferentially consumed over EG. Thus, we prioritized the engineering of TPA metabolism.

Pp_KT2440 naturally degrades protocatechuic acid (PCA)^35^, the product of aerobic TPA degradation pathways^36^. It has been demonstrated that by introducing a 3-step pathway (comprising four enzymes) that converts TPA to PCA, Pp_KT2440 can readily utilize TPA as a carbon source^21,37–39^. Here, we attempted to insert natural or engineered TPA pathways. Specifically, we tested engineered pathways, including the four TPA enzymes from *Comamonas sp.* E6^40^ under strong or weak promoters and ribosomal binding sites, including or not the transporter from the same species or from *Rhodococcus sp.* DK17^41^. We also tested the natural TPA pathway from *P. umsongensis* GO16^31^, under its native regulation (Figure S1). We detected growth on TPA as a carbon source only when the *P. umsongensis* GO16^29,31^ natural pathway, encoded in an operon, was tested.

The heterologous TPA degradation operon, consisting of four enzymes (TphA1, 2, 3, and TphB), one transcriptional regulator (*iclR*), and one transporter (*tphK*) (Figure 2A), was inserted into the Pp_KT2440 genome using the EZ-Tn5 transposon system^42^. Following selection after transformation, a single colony, designated Pp_TPA, was isolated and characterized for its ability to grow on TPA as the sole carbon source in concentrations ranging from 0.08 to 10 mM. Pp_TPA can grow in as low as 0.6 mM TPA (Figure S2). When cultivated in minimal medium with 10 mM TPA, Pp_TPA exhibited a doubling time of 4.1 ± 0.6 hours (Figure 2B). This growth rate is similar to that of naturally occurring TPA-degrading bacteria, *C.* sp. E6 (9.4 hours)^43^, *I. sakaiensis* (8.1 hours)^16^, and *P. umsongensis* GO16 (4.1 hours)^31^; and previously engineered *P. putida* strains (5.2 hours^24^ and 3.5 hours^20^) when tested in similar conditions. Finally, we verified that PCA metabolism in Pp_TPA was not affected (Figure S3).

**Figure 2.**
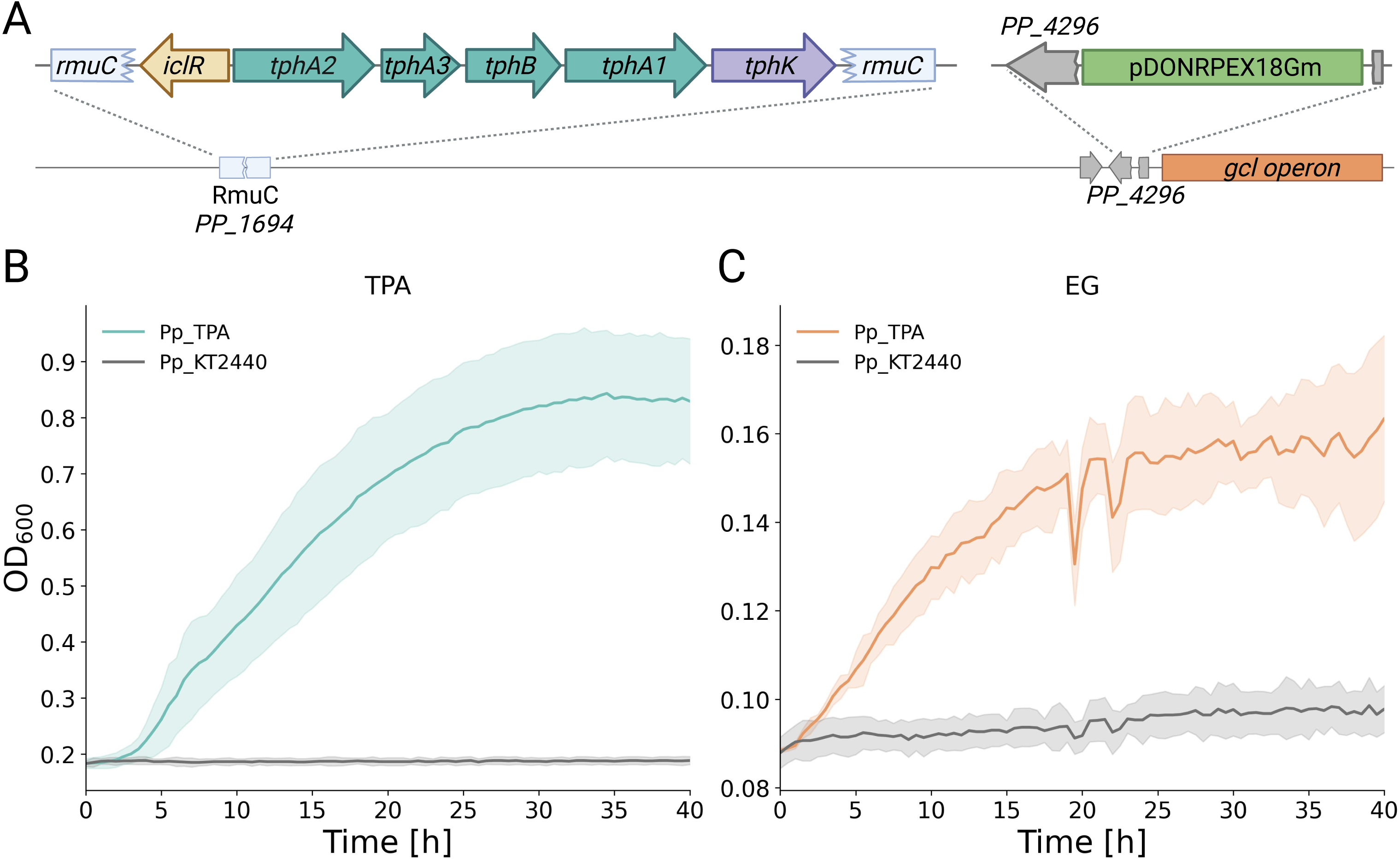
Engineered Pp_KT2440 can use TPA and EG as the sole carbon source. (A) Schematic representation of the Pp_TPA genome in comparison to Pp_KT2440, showing the two identified genetic modifications. One, the insertion of the TPA catabolic operon from *P. umsongensis* GO16 into the *RmuC* gene, and two, the insertion of a knockout plasmid, pDONRPEX18Gm^47^, upstream of the *gcl* operon, disrupting the *PP_4296* gene. (B) Growth curves of Pp_KT2440 (gray) and Pp_TPA (green) in AB medium supplemented with 10 mM TPA. Data represent the mean of 27 growth curves (9 biological replicates with 3 technical replicates each) from 3 independent experiments; shaded areas indicate standard deviation. (C) Growth of Pp_TPA (orange) and Pp_KT2440 (gray) in AB supplemented with 10 mM EG as the sole carbon source. The line represents the mean of three biological replicates with three technical replicates each.

We verified the genomes of Pp_KT2440 and Pp_TPA by sequencing using both Nanopore long reads and Illumina short reads, obtaining high-quality, closed genomes. Comparative genome alignment using Breseq^44^ and Mauve^45^ revealed only two differences (Figure 2A): (1) the TPA pathway is inserted disrupting the *rmuC* gene, which encodes a recombination protein that limits chromosomal inversions, potentially increasing recombination efficiency during pathway integration^46^; and (2) we detected the plasmid pDONRPEX18Gm^47^ integrated upstream to the glyoxylate catabolism (*gcl*) operon, which encodes the enzymes for the last steps of EG utilization. Specifically, the plasmid is disrupting the *PP_4296* gene of unknown function. It originated from previous independent attempts to optimize EG metabolism in our lab. As this unintended genomic rearrangement altered the organization of EG-related genes, we tested Pp_TPA for growth on EG as the sole carbon source. Growth assays showed that, unlike Pp_KT2440, in Pp_TPA, EG metabolism is not silent. Pp_TPA can utilize EG as the sole carbon source in concentrations ranging from 10 mM to 1 M (Figure 2C and Figure S4), indicating that the strain has acquired improved EG metabolism in addition to the introduced TPA pathway.

### Engineering PETases for extracellular PET degradation

After successfully engineering Pp_KT2440 for the intracellular metabolism of PET degradation products (Pp_TPA), we proceeded to find conditions for the extracellular expression of active PETases at 30 °C, the optimal growth temperature for Pp_KT2440. We constructed a library of 14 plasmids expressing either a wild-type or an engineered PETase under the control of different promoters and secretion systems. Two different wild-type enzymes were tested: the leaf-branch compost cutinase (LCC)^48^ and *I. sakaiensis* PETase (IsPETase)^16^, as well as two engineered variants: HOT-PETase^49^ and FAST-PETase^32^. LCC and HOT-PETase, while having optimal temperatures above 30 °C, are highly stable^49,50^, whereas IsPETase and FAST-PETase retain relatively high activity at 30 °C^32,51^.

The four enzymes were fused to the N-terminal signal peptide of the outer membrane porin OprF (SP_OprF_), encoded by the gene *PP_2089* (residues 2-29) of Pp_KT2440. This signal peptide has previously been shown to efficiently export active heterologously expressed enzymes to the extracellular environment^52^. In addition, FAST-PETase was also tested with the signal peptide of the maltotetraose-forming amylase from *P. stutzeri* MO-19 (SP_Pstu_), as used in the original publication where FAST-PETase was described^32^. Besides free diffusion to the extracellular environment mediated by signal peptides, we tested cell-surface anchoring. We chose the EstP protein (PP_0418), an autotransporter of an esterase in Pp_KT2440, which was found to be successful in anchoring heterologous enzymes^52^. The EstP anchor domain (residues 201 to 423) has been attached to the C-terminal of IsPETase or LCC. All constructs, EstP or SP fused, were expressed under weak and strong promoters and RBS. His-tag was added to facilitate purification if needed (Figure S5).

To monitor extracellular PETase activity, we adapted a qualitative halo-based assay using agar plates with BHET and amorphous PET, supplemented with 10 mM glycerol as a carbon source^53^. Since BHET and PET are less soluble than TPA or EG, enzymatic hydrolysis results in localized clearing around the colonies due to loss of turbidity, forming a halo indicative of enzymatic function. Among the tested variants, none expressed with the autotransporter were positive (Figure S6). From the non-anchored variants, we found that all of them were active in plates with BHET when under the control of the weak constitutive promoter, except for HOT_PETase. Under the strong constitutive promoter, only FAST and IS PETases were found to be active (Figure 3B and Figure S7). In the PET plates, we found that only HOT and FAST PETases were active, and exclusively when expressed under the weak promoter and RBS. From the two signal peptides tested in the FAST_PETase constructs, SP_OprF_ was the best solution in all conditions (Figure 3B and Figure S7). Indeed, SP_OprF__FAST-PETase under a weak constitutive promoter reproducibly showed the fastest appearance, clearer, and largest halos in the BHET and PET plates. Hence, it was implemented in the next section.

**Figure 3.**
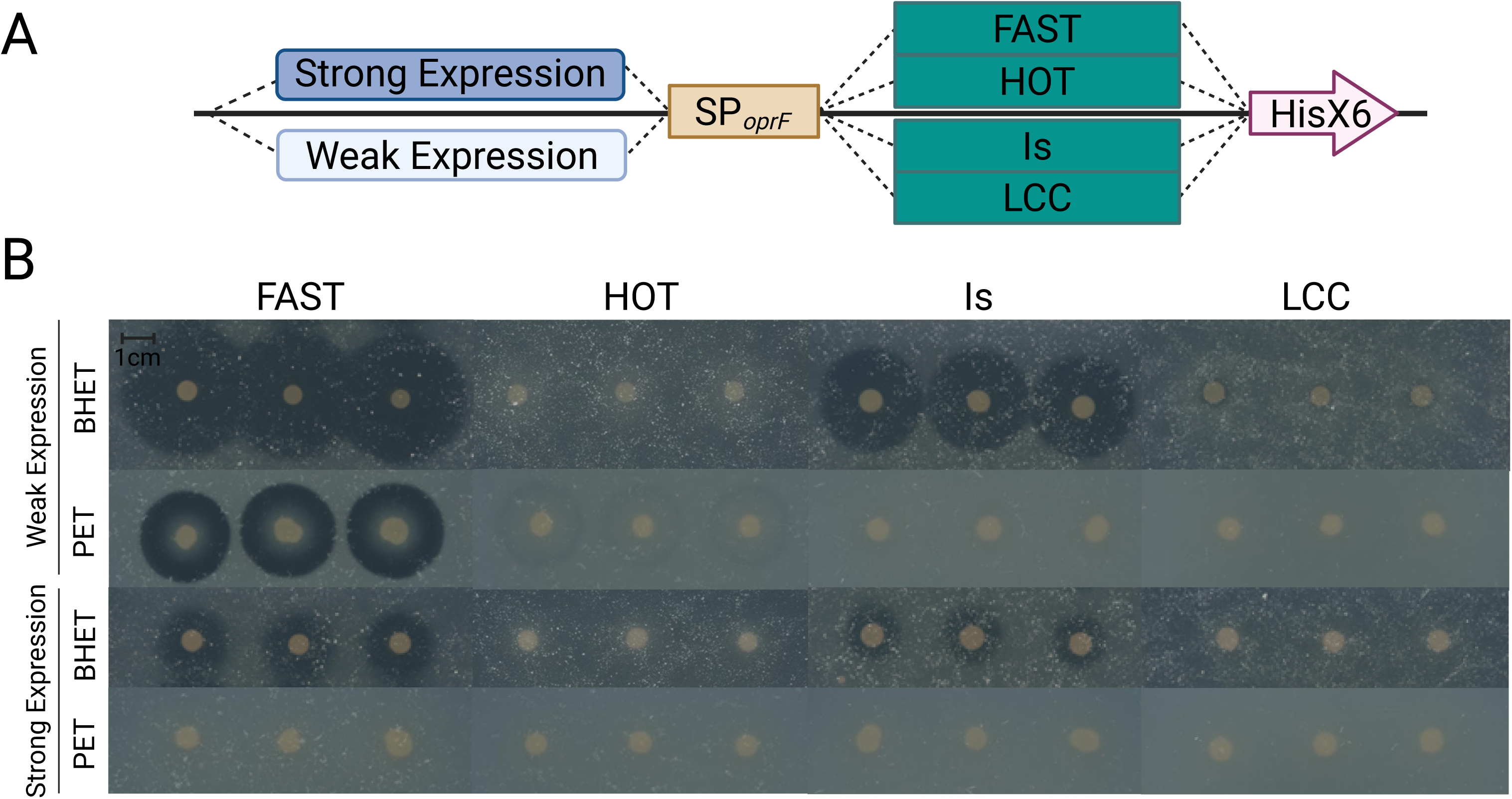
Active extracellular PETases expressed in Pp_KT2440. (A) Schematic representation of the different designs used in this experiment. (B) Images of BHET and PET plates supplemented with 10 mM glycerol and 50 μg/ml Kan. Three colonies represent three technical replicates of each treatment. All pictures were taken after 8 days of incubation.

### Integration of TPA metabolism with PETase expression to achieve growth on PET as a sole carbon source

After successfully developing the essential system components, TPA and EG metabolism, along with the extracellular expression of active PETases, we combined them to establish a complete PET degradation system. Pp_TPA was transformed with the plasmid expressing FAST_PETase under weak expression, secreted via the OprF signal peptide. The resulting strain, Pp_TPA+FAST, was tested for its ability to degrade and assimilate BHET as the sole carbon source in liquid culture (Figure 4A). Indeed, Pp_TPA+FAST exhibited a doubling time of 12.6 ± 3.7 hours in BHET.

**Figure 4.**
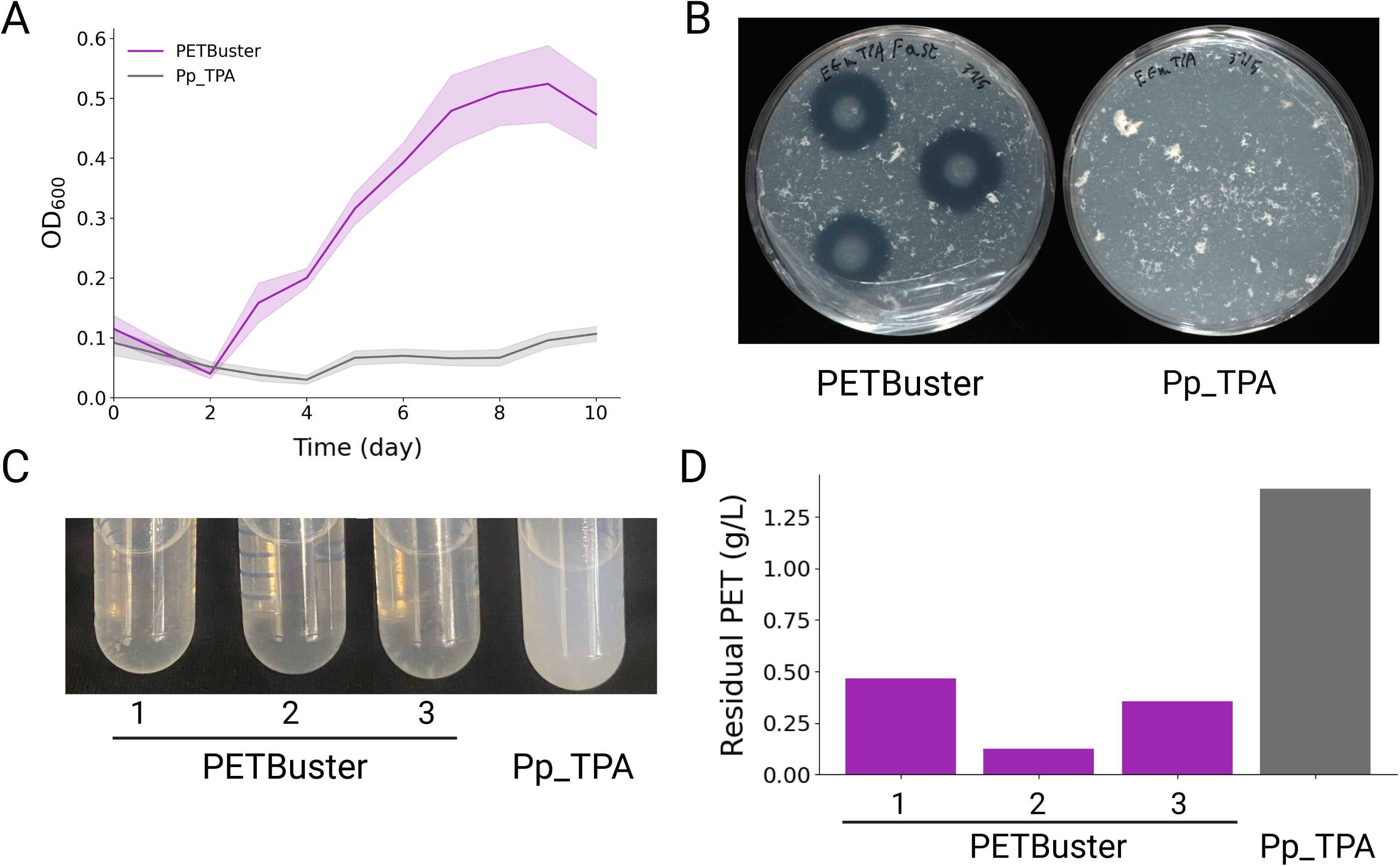
Growth of PETBuster on BHET and PET under different conditions. (A) Growth of PETBuster and Pp_TPA in AB medium supplemented with 10 mM BHET as the sole carbon source. Each curve represents the mean of 12 growth curves from 4 colonies across 2 independent experiments; shaded areas indicate standard deviation. (B) Representative images of PETBuster and Pp_TPA after 20 days of incubation on agar plates containing PET as the sole carbon source. (C) Cultures in AB medium containing pretreated PET after 99 days of incubation. Three tubes were inoculated with PETBuster, and one tube with Pp_TPA as a control. (D) Quantification of residual PET from the cultures shown in panel C, measured by alkaline hydrolysis followed by HPLC analysis (see methods).

We then evaluated Pp_TPA+FAST’s ability to degrade PET and use its degradation products as the sole carbon source under solid and liquid media. Under solid media, we used the same agar plates described in Figure 3, except that the sole carbon source was PET, and no glycerol was added. In liquid, we chemically treated commercially available pure PET powder (>50% crystalline, Goodfellow, Cat. ES30-PD-000132) to reduce crystallinity. This process resulted in less than 10 µM concentrations of BHET, mono-(2-hydroxyethyl)terephthalic acid (MHET, similar to BHET but lacking one EG), or TPA (Figure S8). This concentration cannot support the growth of Pp_TPA. However, Pp_TPA+FAST, hereafter PETBuster (*PolyEthylene Terephthalate Buster*), is capable of breaking down the polymer and utilizing the resulting monomers for growth. Our initial attempts, using 4 mL culture volumes, revealed that after 99 days of culture, biomass accumulation was visible, and 77.3 ± 14.2 % of the PET had degraded (Figure 4C and D).

After initial confirmation that PETBuster could grow on PET as the sole carbon source, we designed a 28-day growth experiment to monitor PET utilization in detail. Six independent PETBuster colonies and six Pp_TPA control colonies were cultured in parallel at 30°C in 30 mL volumes in 150 mL flasks. Bacterial growth was monitored using both colony-forming units (CFU) counts and a resazurin-based viability assay^55^, providing complementary measures of proliferation, metabolic activity, and cell viability. The growth dynamics (Figures 5A and B) indicate that the duplication time of PETBuster, as calculated from CFU measurements, was 3.6 ± 0.8 days (*n* = 6).

**Figure 5.**
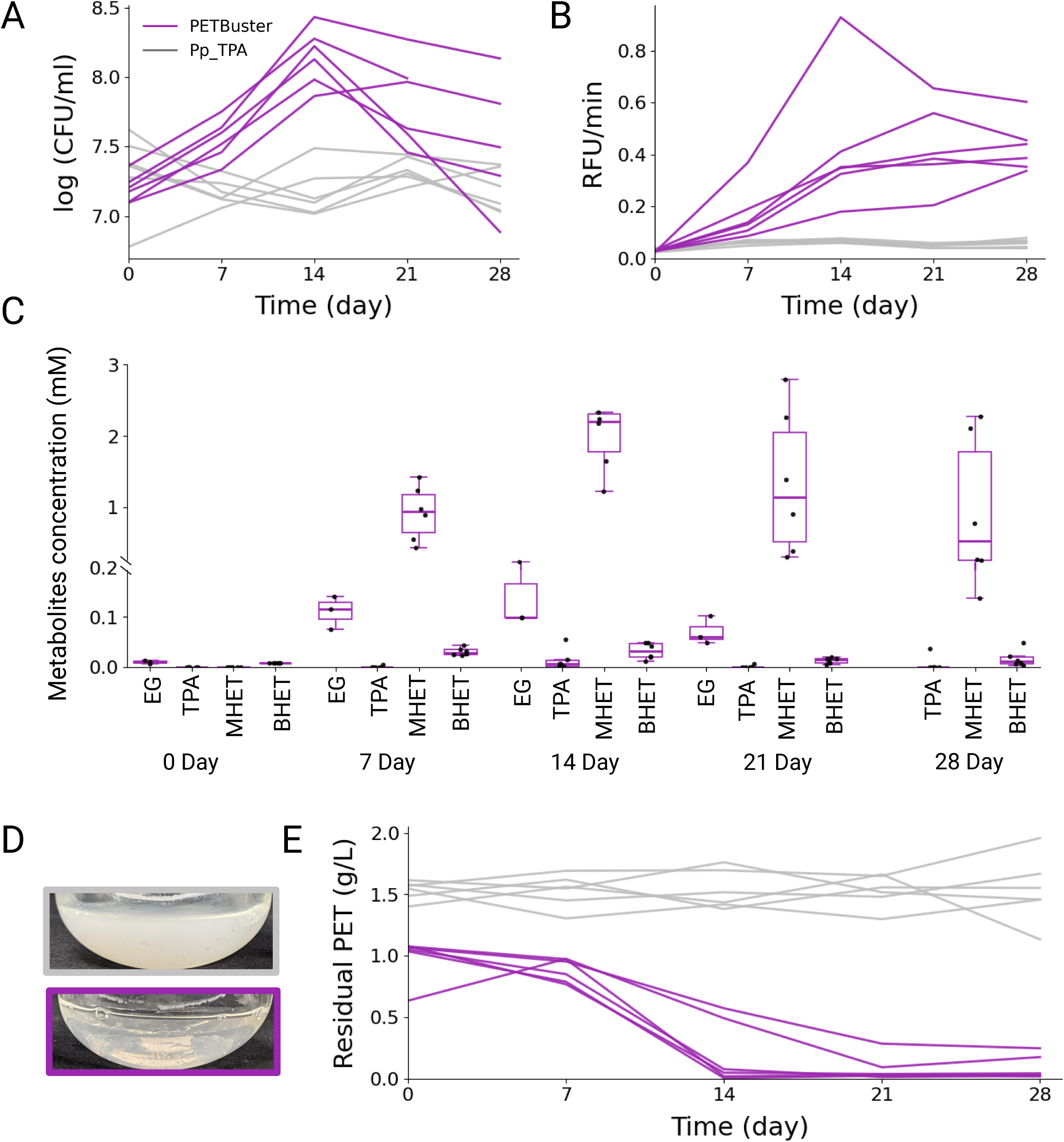
Quantification of PET utilization by PETbuster. (A) CFU counts over time. (B) Resazurin assay measuring cell metabolic activity (viability). (C) Quantification of PET degradation products (EG, TPA, MHET, and BHET) in PETBuster cultures. Boxplots display the distribution of replicate values (n = 6 per condition, n = 3 for EG measurement). The central line marks the median; boxes indicate the interquartile range (IQR); whiskers extend to 1.5 × IQR; and dots represent individual replicates. (D) Representative flask images after 12 days of growth: the upper flask shows Pp_TPA and the lower flask shows PETBuster. (E) Residual PET concentration in the culture during growth.

To evaluate PET utilization over time, we quantified the monomers BHET, MHET, and TPA in the extracellular environment using high-performance liquid chromatography HPLC and EG using gas chromatography - mass spectrometry (GC-MS). All samples were analyzed every seven days (five time points) across the six replicates for all metabolites, except for EG, which was measured in three replicates during the first four time points. We also measure the amount of PET that was not consumed— residual PET (see Methods). As expected, BHET, MHET, and TPA and EG concentrations remain zero or undetectable in the negative control, Pp_TPA, throughout the 28 days (Figure S9). However, in PETBuster growing on PET, we observed higher concentrations of these metabolites at various time points (Figure 5C and S9). TPA and BHET concentrations reached a maximum of 0.06 mM, while those of EG and MHET were 0.25 mM and 3.6 mM, respectively. Indeed, EG and MHET were the slowest to be metabolized, as higher concentrations than TPA and BHET were always detected.

The highest concentration of all metabolites was detected on either day 14 or day 21. After 28 days, all concentrations decreased, indicating intermediates don’t accumulate and are consumed by cells (Figure 5C and S9). After two weeks, it was visible in all PETBuster cultures that turbidity had decreased (Figure 5D). Indeed, residual PET quantifications showed that, after 21 days, PET concentrations had declined by 91 ± 10% (Figure 5D). By contrast, PET levels in control cultures remained stable throughout the experiment (1.5 g/L ± 0.15, *n* = 30). This combined analytical strategy, tracking biomass growth, soluble intermediates, and residual polymer, provides a comprehensive assessment of PET consumption and demonstrates that PETBuster efficiently degrades PET while maintaining robust growth in long-term incubation.

## Discussion

In this study, we successfully engineered *Pseudomonas putida* KT2440 to utilize polyethylene terephthalate (PET) as its sole carbon source. By integrating a heterologous terephthalic acid (TPA) catabolic operon into the genome and extracellularly expressing an active PETase, we created PETBuster, a strain capable of degrading and assimilating PET. When cultured with PET with low crystallinity, the engineered strain degraded up to 91% of the polymer after 21 days, with a doubling time of ∼3.6 days. Importantly, no accumulation of PET monomers (TPA, MHET, or BHET) was detected in the medium, suggesting rapid uptake. Still, we observed that MHET and EG were the slowest to be metabolized, opening the door for further optimization.

Growth was reproducibly observed in multiple independent cultures, as confirmed by both colony-forming unit (CFU) quantification and resazurin-based metabolic activity assays. However, after 2 weeks, we observed a reproducible and significant reduction in biomass. Our HPLC analysis shows molecules with higher molecular weight than BHET. Moreover, our experimental setup, which involves taking a 7% sample volume and long incubation times in flasks, leads to a significant reduction in the working volume and evaporation. These conditions can lead to cell stress, and further investigation is needed to establish the cause of this phenomenon and resolve it. Comparing PETBuster with the three naturally occurring strains that can degrade PET and assimilate its monomers as the sole carbon source, namely, *I. sakaiensis* 201-F6^16^, *R. pyridinivorans* P23^17^, and *P. chengduensis* BC1815^18^, is challenging due to the different conditions used to verify PET metabolism. Particularly, the comparison with *I. sakainensis* 201-F6 is troublesome. When this bacterium was exposed to PET as the sole carbon source, 0.01% yeast extract was in the medium; the contribution of yeast extract to biomass has not been established^16,56^. *R. pyridinivorans* P23^17^ and *P. chengduensis* BC1815^18^ were tested using PET films with more than 50% crystallinity as the sole carbon source. Biomass increased 4 to 22 times, and the PET film lost 1.4% or 4.2% of its weight after one week of cultivation^17,18^.

Similarly, PETBuster biomass increased 5 times; however, we observed 91% PET disappearance. The nature of the PET polymer could be relevant here. It is known that PETases, even highly engineered variants, cannot effectively degrade crystalline PET^57^. The fact that PETBuster was facing amorphous PET (chemically treated with a solvent) could be the reason for the successful degradation of almost all PET in our cultures. It is assumed that enzymatic hydrolysis of crystalline PET would remain elusive. Thus, the most successful approaches for PET biodegradation begin with treatments that reduce crystallinity to less than 10%, allowing for better enzymatic degradation ^54,58,59^, as done here, making PETBuster relevant to industrial applications. Indeed, engineered and optimizable strains, such as PETBuster, open the door to practical microbial strategies for mitigating one of the most abundant pollutants on Earth, while laying the groundwork for future applications in plastic upcycling and sustainable biotechnology.

### Materials and Methods Culturing Conditions

Strains and plasmids used in this study are listed in Table S1. All experiments were performed using either LB or AB minimal medium, as specified. LB was obtained commercially (Difco™). AB medium was prepared following established protocols. Briefly, AB is composed of two solutions: medium A (0.2% (NH₄)₂SO₄, 0.6% Na₂HPO₄, 0.3% KH₂PO₄, and 0.3% MgCl₂) and medium B (0.1 mM CaCl₂, 1 mM MgCl₂, and 0.003 mM FeCl₃). To prevent aggregation, medium A was prepared at 20% of the final volume, and medium B at 80% of the final volume. Component A was prepared and autoclaved, while the liquid stock of Component B was autoclaved separately before mixing. ABx2 medium was prepared at twice this concentration. For all growth experiments, precultures were initiated from single colonies in LB and grown overnight. These precultures were refreshed and washed repeatedly in carbon-free AB medium to remove residual rich medium before being used to inoculate the experimental conditions described.

### Carbon Source Utilization Assays (PCA, TPA, EG, BHET)

Bacterial growth on protocatechuic acid (PCA; Acros Organics, cat. 114891000), terephthalic acid (TPA; Acros Organics, cat. 180725000), ethylene glycol (EG; Sigma-Aldrich, cat. 102466), and bis(2-hydroxyethyl) terephthalate (BHET; Sigma-Aldrich, cat. 465151) was assessed under different conditions. For TPA and EG assays, growth was measured in 96-well microplates using a plate reader (BioTek Synergy H1, Agilent Technologies). TPA was added to medium A prior to autoclaving, then mixed with medium B and adjusted to pH 7.0. TPA was tested at 10 mM and at 7 dilutions, each with a factor of 2. For EG, sterile-filtered stock solutions were added directly to autoclaved AB medium at final concentrations of 1 M, 20 mM, and 10 mM. For PCA assays, a 20 mM PCA stock solution in Milli-Q water was sterile-filtered and diluted with ABx2 medium to yield a final concentration of 10 mM. For BHET assays, growth was monitored in culture tubes (4 mL culture volume) using a Biowave Cell Density Meter CO8000 (Biochrom, catalog no. 80-3000-45). BHET was finely ground with a mortar and pestle before being added directly to sterile AB medium at a final concentration of 10 mM. Cultures were incubated at 30 °C with shaking at 190 rpm, and OD₆₀₀ was recorded at regular intervals. Growth across all assays was monitored by optical density at 600 nm (OD₆₀₀) with continuous shaking at 30 °C unless otherwise specified.

### Growth of PETBuster in solid media

To prepare PET plates with no edible carbon source, 5 g of PET powder (Goodfellow) was dissolved in 15 mL of DMSO by heating the solution on a magnetic stirrer at full power until fully solubilized. Then, the solution was rapidly poured into 1 L of deionized water (DDW) to induce PET precipitation. The suspension was stirred for 1 h, filtered through a 0.45 µm cellulose membrane (Whatman Cat. 10401612), and washed twice with 10 L of DDW. The washed PET was collected, dried at ∼100 °C, and ground into a fine powder using a mortar and pestle. The PET powder (5 g) was suspended in 50 mL of 2x AB medium and sonicated for 6 h (5 s on/off, 50% amplitude) (MRC-Lab, Cat. Sonic-650W) while stirring. The volume was adjusted as necessary, and the PET suspension was combined with 450 mL of additional 2x AB medium and 500 mL of 3% Noble agar. 25 mL of this mixture was poured per plate, yielding PET plates with 5 g/L PET in AB 1x medium and 1.5% agar.

### Growth of PETBuster in PET in liquid media

Cultures were incubated at 30 °C and 190 rpm in AB minimal medium supplemented with pretreated PET -see protocol below- as the sole carbon source (30 mL per flask). Enishel OD 0.025. For inoculum preparation, six biological replicates per strain were initiated from single colonies grown in 4 mL LB, harvested in logarithmic phase, washed four times in carbon-free AB medium, and adjusted to an OD₆₀₀ of 0.025 prior to inoculation. Cultures were incubated for 35 days, with samples collected every 7 days for (i) HPLC analysis of monomer release, including chemical degradation of PET pellets for residual PET quantification, (ii) viable cell counts by CFU plating on LB agar, and (iii) metabolic activity measurements using a resazurin-based fluorescence assay.

### Measuring growth with resazurin

Redox metabolic activity was assessed using a resazurin reduction assay adapted for a 96-well microplate format. Each reaction well contained 90 μL of bacterial culture and 10 μL of a freshly prepared resazurin solution (final concentration: 0.01 mg/mL).

Reactions were incubated in a Synergy H1 microplate reader (BioTek Technologies) maintained at 30°C, and fluorescence was monitored every 4 minutes for 48 hours with constant shaking. Fluorescence readings were collected from the bottom using excitation at 535 nm and emission at 590 nm. The raw fluorescence units (RFU) were analyzed by calculating the maximum slope of RFU over the first ⅓ of the reds using 10 data points over time to estimate resazurin reduction rates.

### Measuring growth with Colony Forming Unit (CFU)

Bacterial growth was quantified by CFU counting using serial dilutions. From the 10⁻³ and 10⁻⁶ dilutions, 100 µL was plated on LB agar plates and incubated overnight at 30 °C. Colonies were counted the following day, and CFU/mL was calculated by multiplying the colony number by the reciprocal of the dilution factor and an additional factor of 10 to account for plating 100 µL.

### Calculations of the duplication times

For each replicate growth curve, optical density data were log-transformed, and the specific growth rate (µ h⁻¹) was obtained using a sliding-window linear regression: a 2 h window for TPA growth experiments and a 24 h window for BHET growth experiments. Minimal duplication times were calculated as ln(2)/µ. For CFU-based measurements, duplication time was estimated from the change in viable counts between days 7 and 14. Duplication times from previously published studies were extracted manually by selecting two data points from published growth plots using a web tool, and calculating µ and doubling time as above.

### Plasmid Construction

All PETase sequences were ordered from Twist Bioscience. The signal peptide of oprF (SPOprF, gene PP_2089; residues 2–29) and the EstP autotransporter domain (PP_0418; residues 201–423) were amplified from the P. putida KT2440 genome. The TPA degradation pathway and two TPA transporters were also synthesized by Twist and subsequently cloned using the primers listed in Table S2.

PETase constructs carrying SPOprF or EstP signal peptides were assembled into the pSEVAb237 backbone (low expression by default), while TPA pathway constructs were cloned into the pSEVAb227 backbone from the SEVA plasmid collection^60^. DNA assembly was performed using the NEBuilder HiFi DNA Assembly kit (New England Biolabs, NEB). PCR amplifications were carried out using either KAPA HiFi HotStart ReadyMix or Q5 High-Fidelity 2X Master Mix (NEB). PCR products were verified by agarose gel electrophoresis.

Assembled plasmids were verified by sequencing (either full plasmid or insert only) using Oxford Nanopore or Sanger sequencing. For cloning steps, chemically competent *E. coli* TOP10 cells prepared in-house were used. Final plasmids were introduced into *P. putida* KT2440 using a standard electroporation protocol^61^.

### Engineering the Pp_TPA strain

We received the TPA pathway in the plasmid pBT’T^38^ from Pablo Nkel Lab at DTU Biosustain. For the insertion of the TPA pathway into the Pp_KT2440 strain genome, we used the EZ-Tn5 transposon kit (Biosearch, Cat. No. TNP92110), which utilizes the transposase mechanism to randomly insert DNA into the genome. We followed the manufacturer’s conditions with a few modifications. In the transformation reaction, we added 1 µL of TypeOne Restriction Inhibitor from Biosearch Technologies (Cat. No. TY0261H). We used growth on TPA as the selection method for positive strains. After transforming the transposomes complex, the cells were recovered in SOC media (Formedium and Cat. SOC0201) at 30°C with shaking for 2 hours. Then, the cells were centrifuged at 4000 rpm for 3 minutes. We resuspended the cells in AB + 10 mM TPA at pH 7 and left them to grow for 2 days. After 2 days, the culture reached OD 1.35. We transferred 250 µL of the first culture to a fresh 5 mL of AB + 10 mM TPA at pH 7. The second culture reached OD 1.3 after 3 days of growth. We isolated and tested the growth of 28 colonies. Colony 14 showed the best growth on TPA, so we renamed it Pp_TPA.

### Sequencing and comparative analysis of Pp_TPA and Pp_KT2440

Whole-genome sequencing was performed for *Pseudomonas putida* KT2440 and its engineered derivative *P. putida* Pp_TPA by Plasmidsaurus using a hybrid approach with Oxford Nanopore long reads and Illumina short reads. Hybrid assemblies were generated by assembling the long reads and subsequently polishing with the short reads to improve base accuracy. The final polished assemblies were provided by the sequencing service for downstream analyses. Comparative genomics between the two strains was carried out using Mauve (progressiveMauve algorithm, default parameters) to identify structural rearrangements and Locally Collinear Blocks. In addition, single-nucleotide variants (SNVs) and small insertions/deletions were assessed using *breseq* (default consensus mode), with Illumina reads of Pp_TPA aligned against the polished Pp_KT2440 of our lab. This analysis revealed no sequence differences between the two strains.

### Screening PETase function in plates containing BHET/PET with Glycerol

We prepared the AB media with 10 mM glycerol, 10 mM BHET, and 1.5% noble agar. All the compounds were sterilized by autoclave (120°C, 20 min), except BHET, which was treated at 70°C for 1 hour due to its thermo-sensitivity. For the preparation of the PET plate, we used PET powder (with>50% crystallinity, Goodfellow Cat. ES30-PD-000132). We dissolved the PET in DMSO (200 mg of PET into 8 mL of DMSO) and then added it to 80 mL of media.

