## Supplementary Figures for "A synthetic bacterium that degrades and assimilates poly(ethylene terephthalate)"

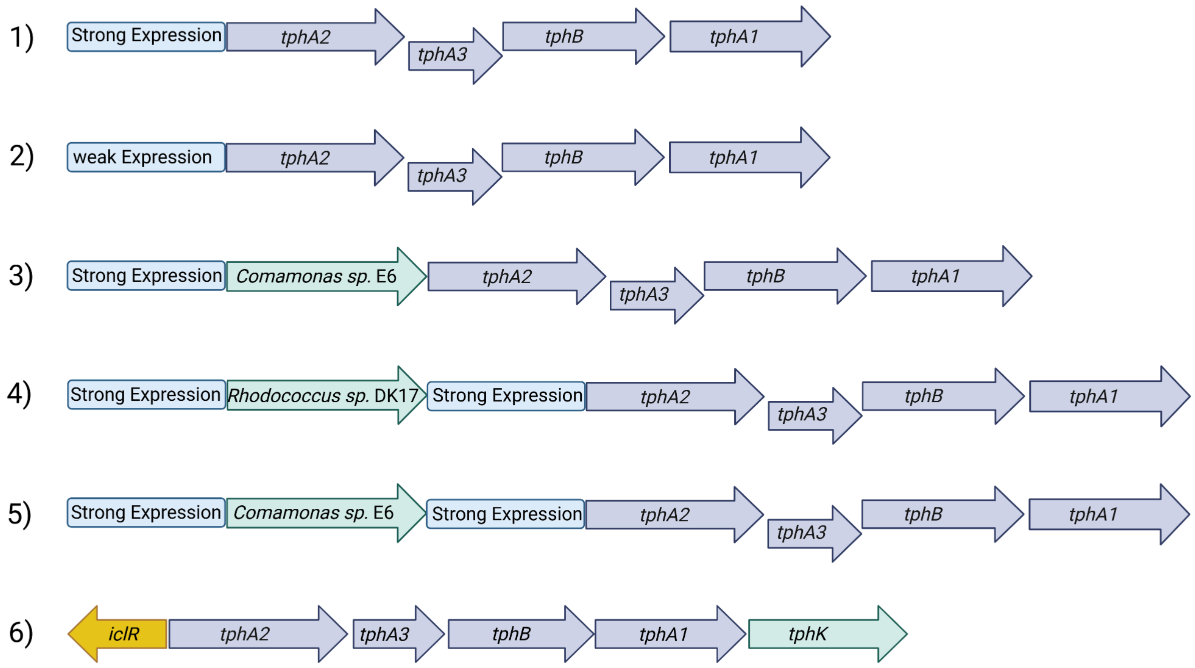


**Figure S1. TPA utilization pathways that were tested in this study.** Schematic representation of the six TPA-utilization constructs generated in Pp_KT2440. The constructs were assembled using different combinations of promoter–RBS expression modules: a weak module (BBa_J23105 + RBS20) or a strong module (BBa_J23100 + RBS15)[3]. Transporters were sourced from *Comamonas* sp. E6[9] or *Rhodococcus* sp. DK17[10]. The TPA catabolic modules included either the degradation pathway from *Comamonas* sp. E6[9] or the TPA degradation operon from *Pseudomonas umsongensis* GO16[8,11]. Constructs 1–5 were built into the plasmid backbone pSEVAb237[3], while construct 6 was cloned from the pBT'T plasmid[2] and integrated into the genome of Pp_KT2440 using the EZ-Tn5 transposon system[12].

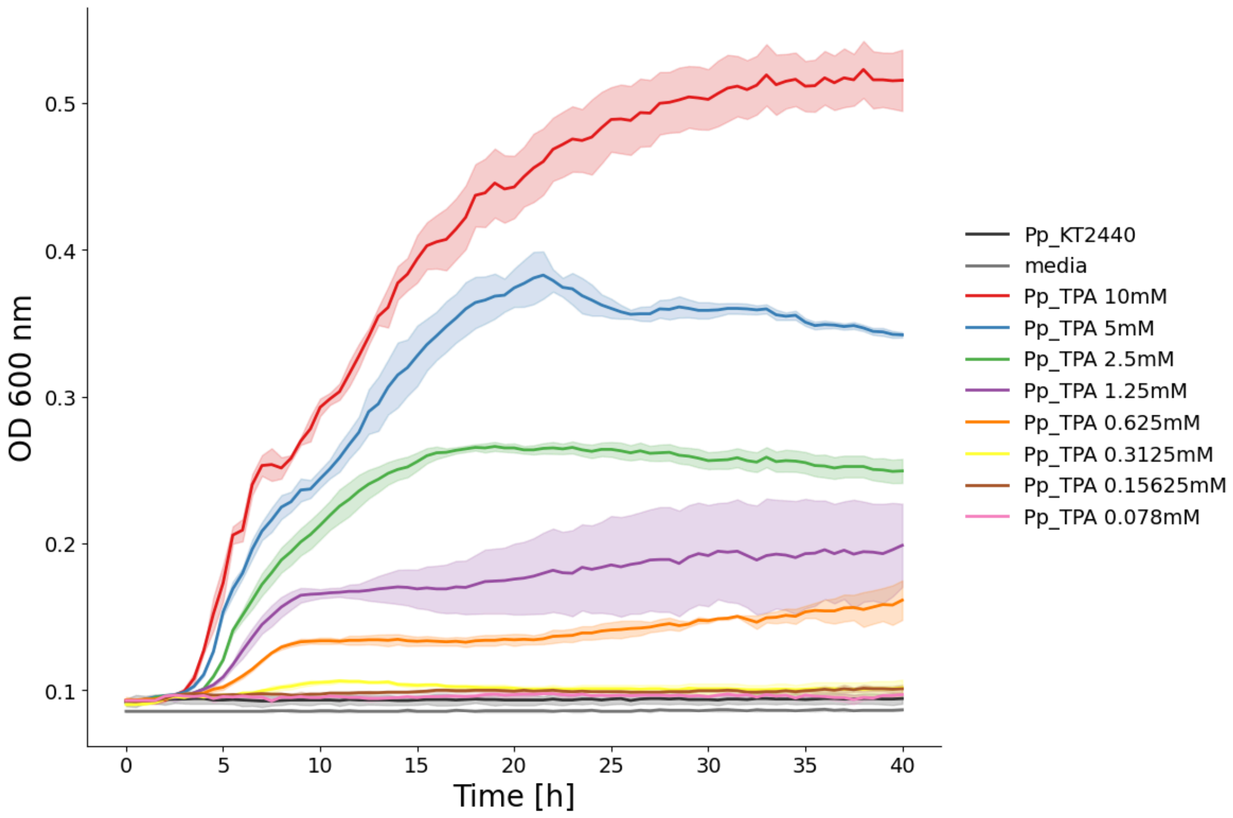


**Figure S2. Minimal TPA concentration supporting the growth of Pp_TPA.** Growth of Pp_TPA was assessed in eight different TPA concentrations ranging from 0.078 to 10 mM. As controls, Pp_KT2440 was grown in 10 mM TPA (black), and AB medium without bacteria is shown in grey. Each line represents the mean of three technical replicates, and the shaded areas represent the standard deviation.

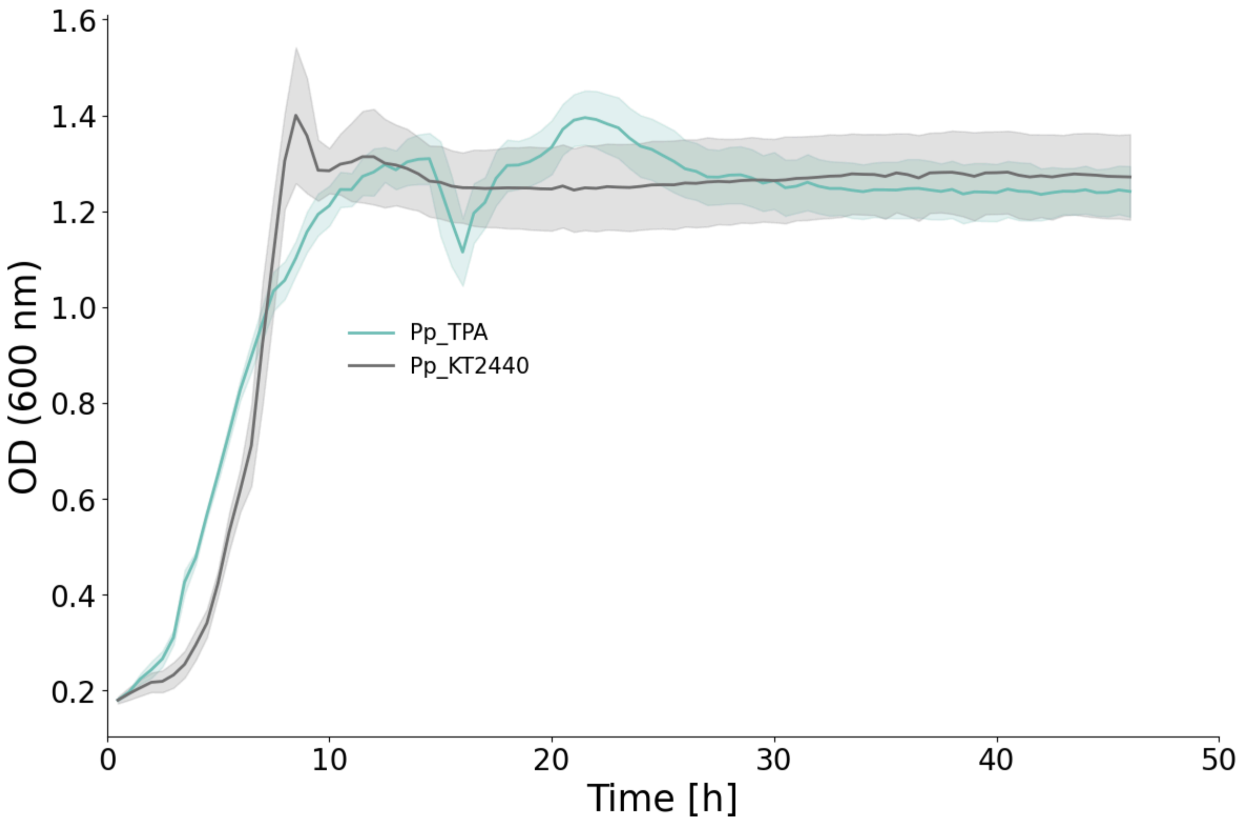


**Figure S3. Growth of Pp_TPA and Pp_KT2440 on PCA.** Growth curves of Pp_TPA (green) and Pp_KT2440 (gray) in AB medium supplemented with 10 mM protocatechuic acid (PCA). Each curve represents the mean of 9 replicates (derived from three independent colonies, with three technical replicates each), and the shaded areas indicate the standard deviation.

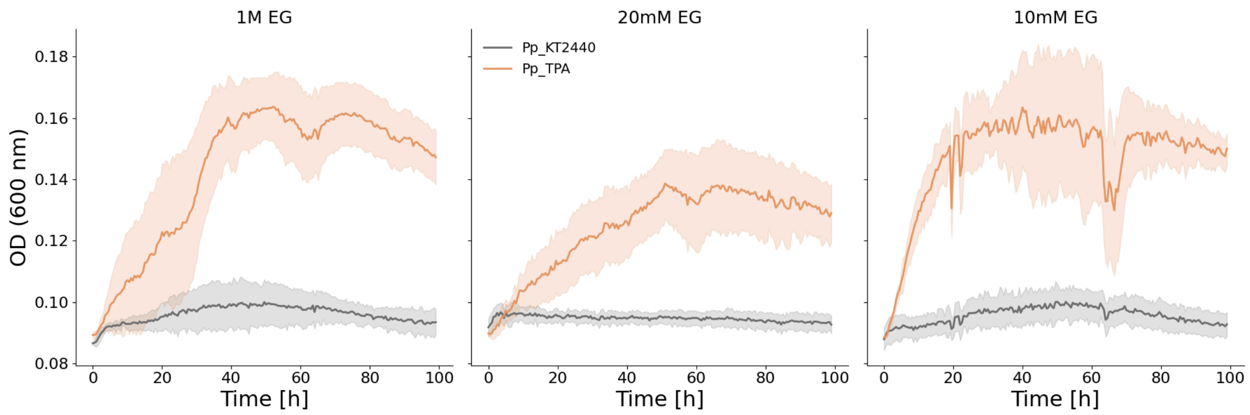


**Figure S4. Growth of Pp_TPA on two different concentrations of EG.** Growth of Pp_TPA on EG as the sole carbon source. The three plots represent growth at different EG concentrations: 1 M, 20 mM, and 10 mM. The Pp_TPA strain is shown in orange, and Pp_KT2440 in grey. Each line represents the mean growth of three biological replicates (independent colonies), each measured in triplicate (three technical replicates per colony). Optical density at 600 nm was measured every 30 minutes. The experiment was conducted in a 96-well plate at 30 °C with continuous shaking.

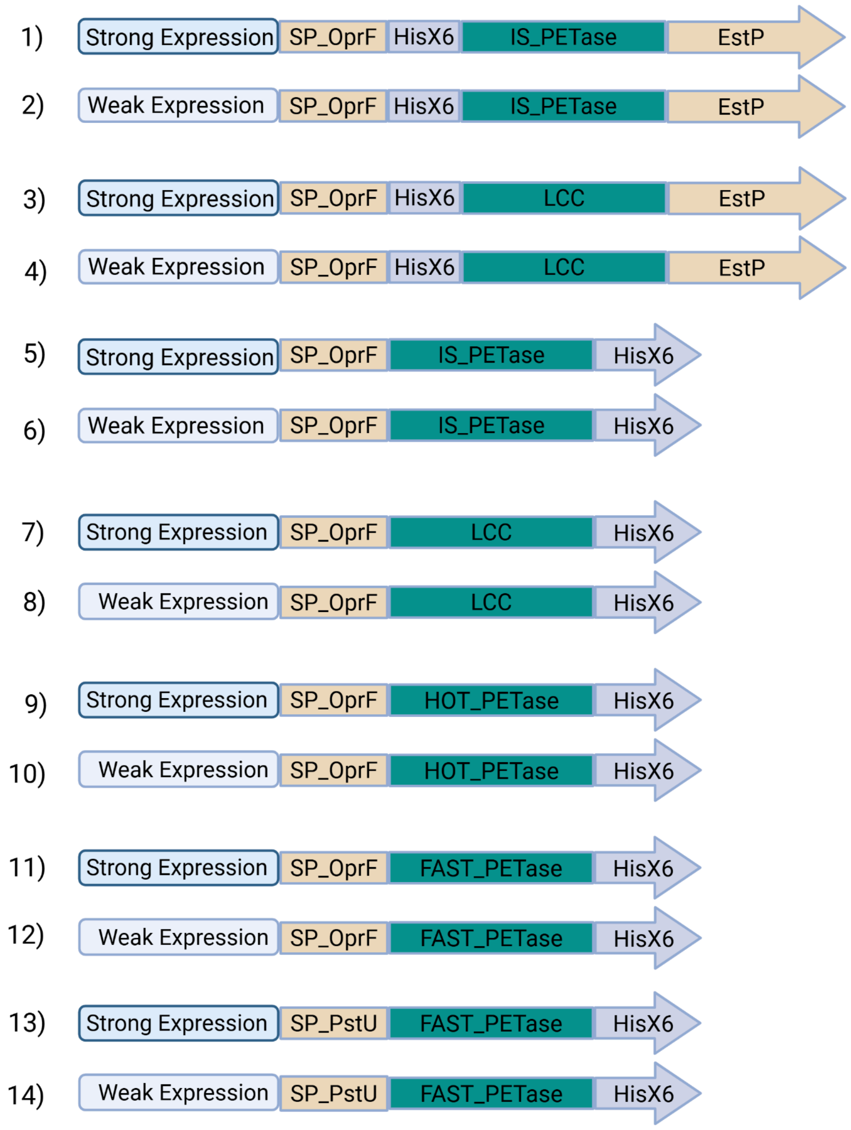


**Figure S5. Constructs for the heterologous expression of PETases in Pp_KT2440.** Schematic representation of the 14 constructs generated in this study for PETase extracellular expression. The library was assembled from nine functional parts: a weak or strong expression module comprising a promoter and RBS combination. The weak expression module has the BBa_J23105 promoter and the RBS203. The strong expression module has the BBa_J23100 promoter and the RBS15[3]. A signal peptide from the *oprF* gene (amino acids 2 to 29)[13], one of four PETases (Is, LCC, HOT, or FAST-PETase)[4–7], the anchoring domain from the EstP protein (amino acids 201 to 423)[13], and a His-tag. The information on all parts is presented in Table S1.

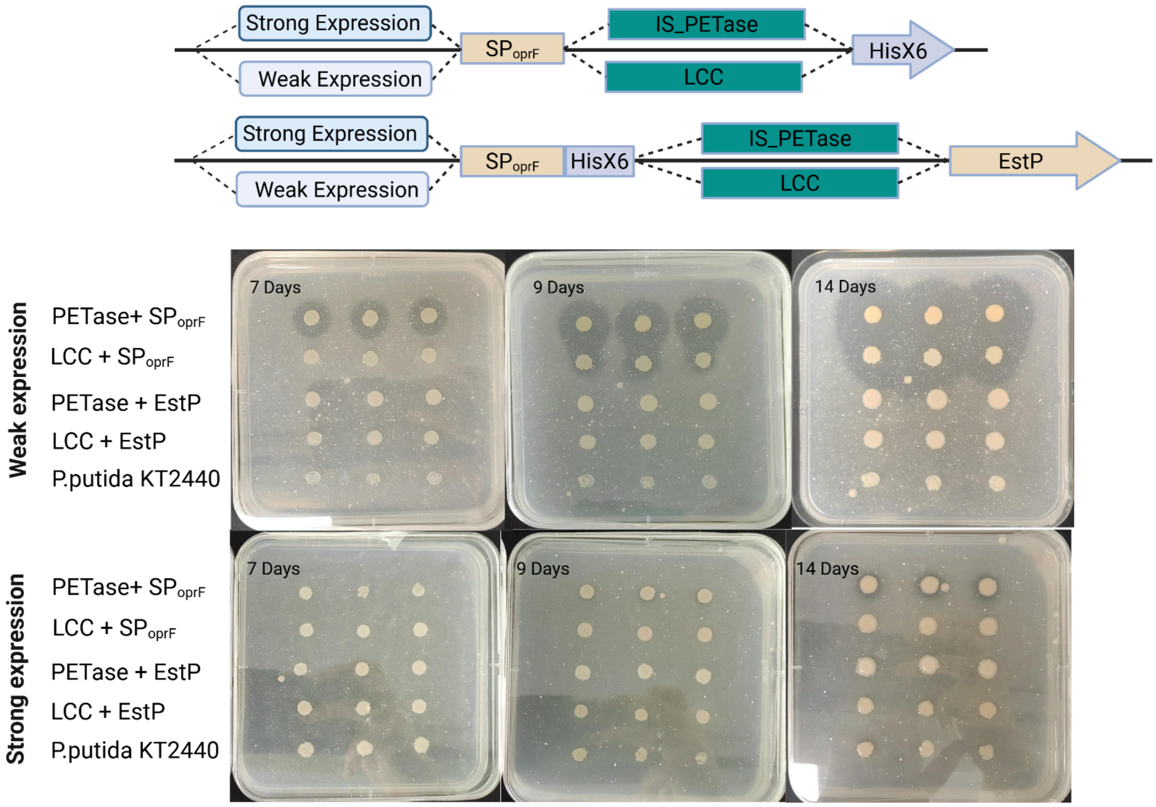


**Figure S6. PETase secretion and activity tests using EstP and SP$_{OprF}$.** Initial experiments monitoring the activity of LCC and IsPETase under weak and strong expression conditions are shown. Secretion was tested through SP$_{OprF}$ or with the EstP autotransporter. PETase activity is indicated by the appearance of halos. Each row shows three independent colonies as biological replicates. Images were captured at three time points: 7, 9, and 14 days.

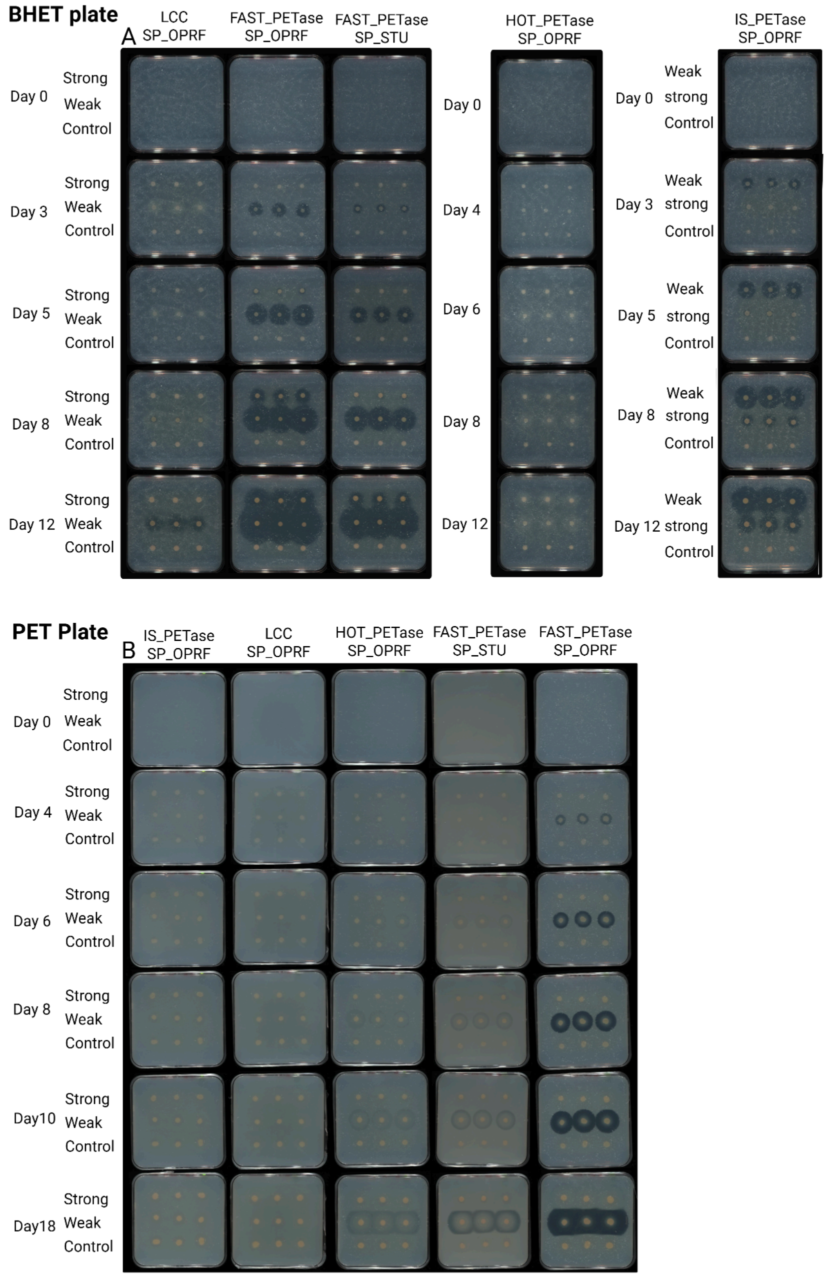


**Figure S7. Halo assay comparing extracellular PETase activity across expression levels and signal peptides.** Colonies expressing each PETase variant were plated on agar supplemented with either (A) BHET or (B) PET. Each plate displays three rows per variant: strong expression, weak expression, and wild-type control (no PETase). For FAST-PETase, two signal peptides—SP_OprF and SP_PstU—were evaluated for secretion efficiency. Formation and size of halos around colonies indicate relative extracellular PETase activity.

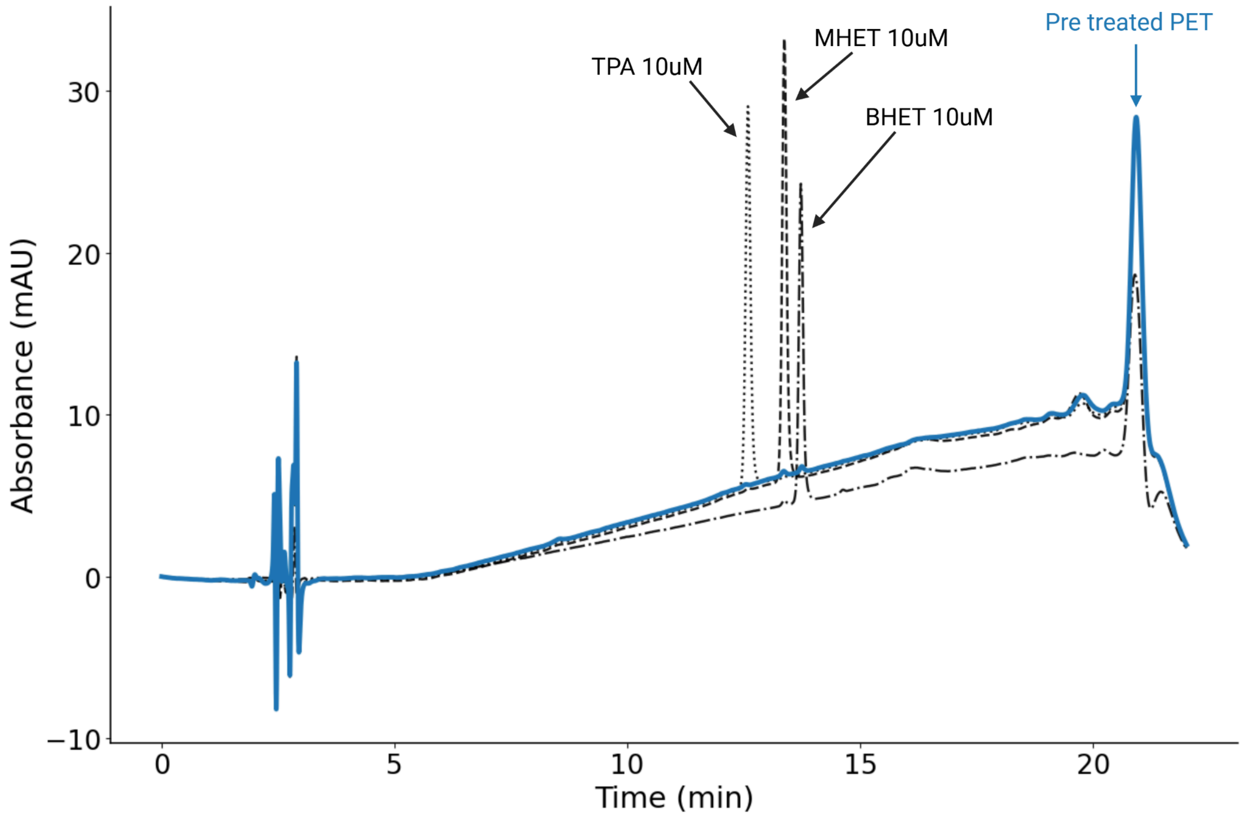


**Figure S8. PET monomer profile following PET pretreatment.** HPLC chromatogram (wavelength 240nm) of the PET supernatant compared with reference chromatograms of 10 µM TPA, MHET, and BHET (shown as dashed lines in blue). The comparison indicates that the concentrations of PET-derived monomers released into the medium after pretreatment are below 10 µM. As shown in Figure S2, such concentrations are insufficient to support Pp_TPA growth.

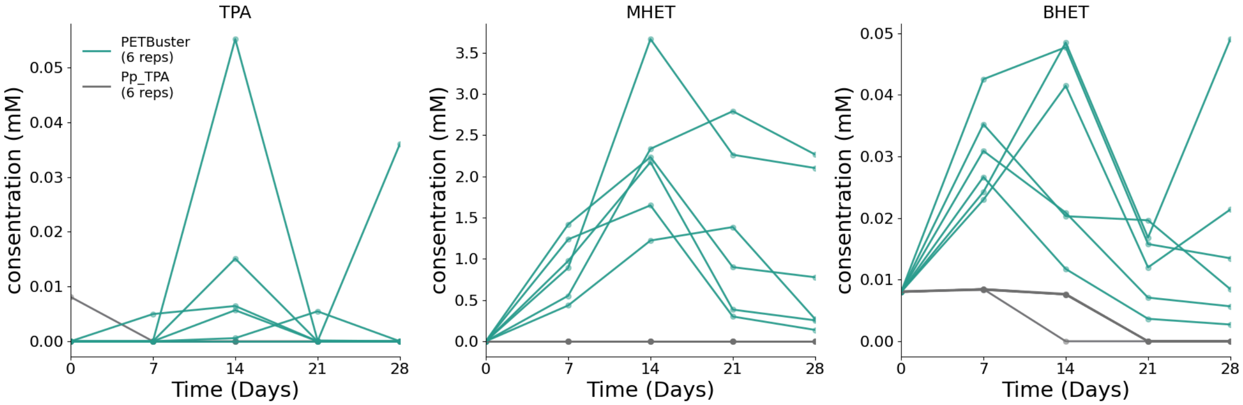


**Figure S9. PET degradation products are formed during growth on PET as the sole carbon source**. Concentration of PET degradation products (TPA, MHET, BHET) during 28 days of growth on PET as the sole carbon source. Each line represents an independent culture (from a single colony), showing the accumulation dynamics of individual degradation products over time.

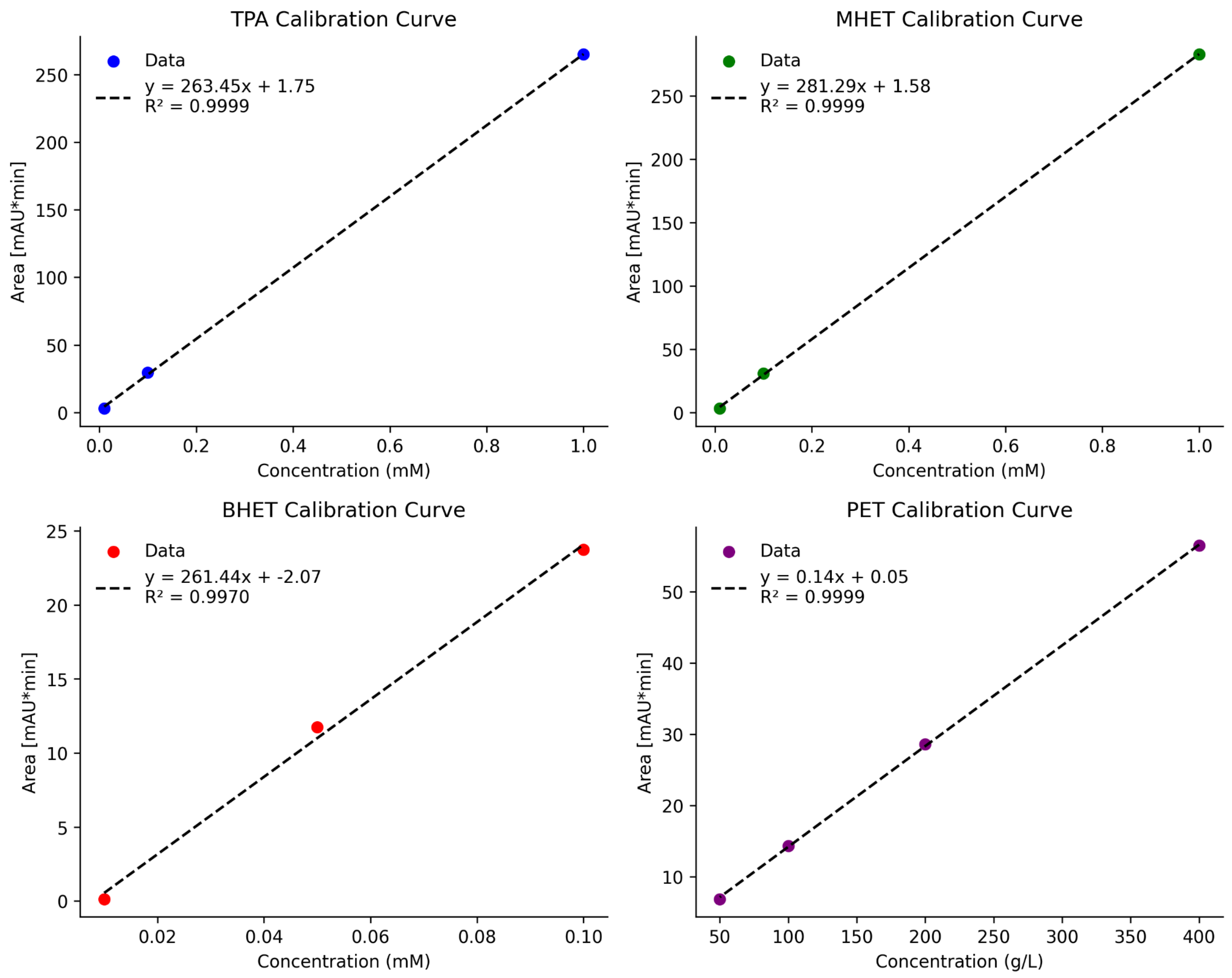


**Figure S10. HPLC calibration curves for PET monomers and polymer.** Peak area (mAU·min) is plotted against standard concentration for TPA, MHET, BHET, and PET. Concentrations are in mM for TPA, MHET, and BHET, and in $g \cdot L^{-1}$ for PET. Points represent injected standards. Dashed lines are least-squares linear fits; the fit equations and $R^2$ values are shown on each panel.
